# The Hidden Repertoire of Brain Dynamics and Dysfunction

**DOI:** 10.1101/578443

**Authors:** Anthony R. McIntosh, Viktor K. Jirsa

**Affiliations:** Rotman Research Institute, Baycrest, Univ of Toronto, Toronto, Canada; Institut de Neurosciences des Systemes, INSERM, Aix-Marseille Universite, 13005 Marseille, France

**Keywords:** Dynamical systems, epilepsy, cognition, neuroimaging, computational modeling

## Abstract

The purpose of this paper is to describe a framework for the understanding of rules that govern how neural system dynamics are coordinated to produce behavior. The framework, structured flows on manifolds (SFM), posits that neural processes are flows depicting system interactions that occur on relatively low-dimension manifolds, which constrain possible functional configurations. Although this is a general framework, we focus on the application to brain disorders. We first explain the Epileptor, a phenomenological computational model showing fast and slow dynamics, but also a hidden repertoire whose expression is similar to refractory status epilepticus. We suggest that epilepsy represents an innate brain state whose potential may be realized only under certain circumstances. Conversely, deficits from damage or disease processes, such as stroke or dementia, may reflect both the disease process per se and the adaptation of the brain. SFM uniquely captures both scenarios.

The Hidden Repertoire of Brain Dynamics and Dysfunction

## 1. Vignette

Both of us like to run, partly for fitness and partly for mental health. It’s easy and you can do it almost anytime and anywhere. The thing about running is that the rules on how to do it are fairly simple, but how you do it is quite varied. Running in the heat of the summer on a beach is different from running up a hill in the forest or trying to navigate an icy trail in the winter. The point here is that while the rules for running are always the same, you would not assume that the example of running on beach serves as an accurate characterization of all running that we might do. The analogy is meant to suggest this is the approach that we use when trying to link brain and behavior. The coordination of behavior by the brain can be understood as a reflection of general rules whose specific realization depends on the current context and initial conditions.

Stated more boldly, we often assume that the expression of behavior at a point in time is sufficient to understand how that behavior is coordinated. Experimental approaches focus on the characterization of brain signal time series and how they change with manipulation. Theoretical approaches most often focus on defining functions that generate these time series. Such approaches are valid insofar as they able to characterize the local conditions that generate the time series. If the nervous system of study can only generate that time series, then this approach will be successful.

However, a different scenario emerges when we consider that a given realization is but one of many that the brain can generate. The brain is a complex adaptive system, showing the properties of multiscale behavior, emergence and nonlinearity(Fingelkurts, 2004; Mitchell, 2009). If we acknowledge this, then a single realization captures only a partial picture of what is possible. Changes to the initial conditions for generation of the behavior can change the realization to the point where the time series bears little resemblance to other realizations. This would be construed as “noise” in most perspectives, but the case we wish to make here is that such variations can be considered a valid expressions of the rules under which behavior is coordinated.

This perspective can be more saliently appreciated when we consider clinical conditions and the variation in expression across persons. For instance, in the case of focal damage from stroke, two persons can show similar regional damage, yet show quite different clinical outcomes (Price & Friston, 2002). Person A may be very impaired, whereas Person B shows remarkable recovery. Person B, in our framework, is less debilitated because they have more options to realize a particular behavior than Person A. The rules that govern behavior are effectively the same for both persons, but the variation in expression is greater in Person B. The stroke impairs one particular set of realizations (i.e., a specific trajectory) abolishing the behavior in Person A, but for Person B only slightly alters the execution. The differences are often explained as resilience or brain reserve, which merely relabels the outcome rather than providing a mechanism of explanation. We propose these mechanisms can be captured in the Structured Flows on Manifolds (SFM) framework (Pillai & Jirsa, 2017).

## 2. General Perspective

We present a framework wherein complex brain dynamics can be decomposed into probabilistic functional modes. These modes are mathematically operationalized as manifolds, along which trajectories evolve as the dynamics unfold embedded in a low-dimensional space or SFM (Huys, Perdikis, & Jirsa, 2014). The collection of functional modes available in a neural network constitutes its functional repertoire, which together instantiates a complete set of potential cognitive functions and overt behaviors.

It has been acknowledged by a number of neuroscience researchers that the brain is dynamic, but how that translates to their approach to gain understanding varies widely. At one end, some consider the brain to be simple input-output system where a signal comes in, a cascade is triggered as the signal propagates, and the system produces an output appropriate to the input (Petersen & Fiez, 1993; Posner, Petersen, Fox, & Raichle, 1988). Other perspectives, stemming from the focus on intrinsic activity in the brain, goes from a unidirectional input-output system to one where the input signal itself may be modified (Deco, Jirsa, & McIntosh, 2013; Fox et al., 2005; Raichle, 2010). One expression, which falls under general category of *predictive coding*, focuses on the time series of neural signals as manifestations of internal models that the brain generates to predict its inputs and its ultimate consequences (Friston, 2010; Rao & Ballard, 1999). There is another elaboration of this that reflects our SFM framework, which we will cover shortly.

The assumption underlying much predictive coding work is that the expression of behavior at a point in time is sufficient to understand how that behavior is coordinated. Other theoretical approaches focus on defining mathematical functions for behavioral time series, while empirical studies use machine learning algorithms to classify the time series according to the behavior they are thought to support (e.g., perceptual categorization).

There are two challenges here. First, if we were to reverse engineer a system that produces the observed time series that reflects the behavior of interest, we would not learn how the behavior itself was coordinated. Rather we would only know what generates individual time series (e.g., the action of a specific set of brain areas). Second, and more problematic, is that the model would not be able to generate new behaviors that we had not previously measured. Said differently, we may be able to predict what the system *has done*, but cannot predict what it *will do*. One remedy is to update the model in light of the new behavior and building a lookup table that relates the configuration of neural dynamics to a specific behavior. The process continues until at some point we have cataloged all the behaviors of the system. While this sounds cumbersome, you see it played out in modern neuroscience. In neuroimaging, for example, we started with the characterization of activated brain regions and relating that to specific behavioral functions (vision, audition, language, memory), drawing inferences on the unobservable processes that were needed to instantiate such functions. We are now in the era of brain networks, where the coherent interactions between regions are the substrate for function (default network, salience network, dorsal attention network). A great deal of research now emphasizes the system characteristics that support these networks by looking a graph theory metrics (Bullmore & Sporns, 2009; Rubinov & Sporns, 2010) and by characterizing feature of the dynamics, such as scale-free behavior and criticality (Beggs & Plenz, 2003; Haimovici, Tagliazucchi, Balenzuela, & Chialvo, 2013; Petermann et al., 2009; Tagliazucchi, Balenzuela, Fraiman, & Chialvo, 2012). If we pause and examine these observations, we have indeed done a good job of characterizing *what* the system does, but have no idea *how and why*.

The SFM framework takes a different approach, which still assumes that the brain constructs models of the world (as in predictive coding), but takes the focus from the specific instantiation of that model (aka the individual trajectory) to discovering the rules that the brain uses to develop these models. This is a subtle, but critical, difference. As we describe the option that encapsulates SFM, it is useful to borrow an analogy from J.H. Holland on the game of chess to illustrate the difference (Holland, 2014). One can learn chess by watching a game and tracking the movements of each piece, repeating the observation for subsequent games and then build a catalogue of moves and counter moves. This is a formidable challenge given that, by rough calculations, there are at least 10^50 possible legal move sequences, which is larger than the estimated number of atoms in the universe. The more efficient approach is to define the rules that determine the legal moves. By doing this for chess, we dramatically reduce the problem from an essentially infinite space to one where a dozen or so rules capture all possible realizations of the chess game. Mastery of chess is achieved when individual moves are combined and orchestrated into larger motifs, further classified into aggressive, defensive and strategic patterns. We understand chess by understanding the rules of play, and understand it deeply by using these rules to build coordination motifs. And this is the option that motivates the description of SFMs: *the goal for understanding brain and behavior is to determine the rules that govern the coordination of behavior*.

Another illustration that makes the distinction between the emphasis on a specific realization versus a model for the rules that generate the realization comes from an example of calculating 3 times 4, 3 time 5, and then switch to 13 times 14, in which the majority of people will rapidly access their semantic memory for the first two cases, but evoke a different model to compute algorithmically the last. If the result of the computation is not in memory, then no solution can be found, whereas in the algorithmic case solutions for number computations may be found that have never been computed before. The most innate and pertinent characteristic of the brain is its capacity to generate dynamic models.

## 3. Conceptual Description of Structured Flows on Manifolds

The SFM framework lies firmly in the ideas of complex adaptive systems (CAS). Our exposition will thus borrow heavily from analogies of other, non-neural, systems that illustrate key principles to build our case, such as emergence, nonlinearity, motifs, flows and internal models.

The notion of SFM formalizes some key general properties of CAS. The use of the term *flows* in SFM emphasizes the dynamic nature of brain processes, where the flow formalizes the rules that enact the *internal model* of the system. The *nonlinearities* of the system impart other properties, such as *aggregation* and *emergence* that link the actions at one level of the system (e.g., network dynamics) to actions at another (e.g., behavior). The elements (or more often called “agents”) can operate at different *timescales*, and the interactions between scales are a critical feature in controlling the flow of the system. Fast time scales may have no overt consequence until slower moving scales reach a certain *tipping point*, or *bifurcation*, and the flow of the entire system changes.

SFM approaches emphasize the manifolds that can be understood as force fields generating the ensemble of all possible trajectories (or flows), and are thus a mathematical expression of the rules underlying the generation of behavior. Figure 1 demonstrates the general idea of flows on manifolds with one toy example of a spherical manifold having two attractor states or domains that support different flows. The figure also demonstrates a comparable manifold architecture in simulated resting stating functional MRI data, where changes in functional connectivity dynamics (FCD) switch two states that span a manifold.

**Figure 1.**
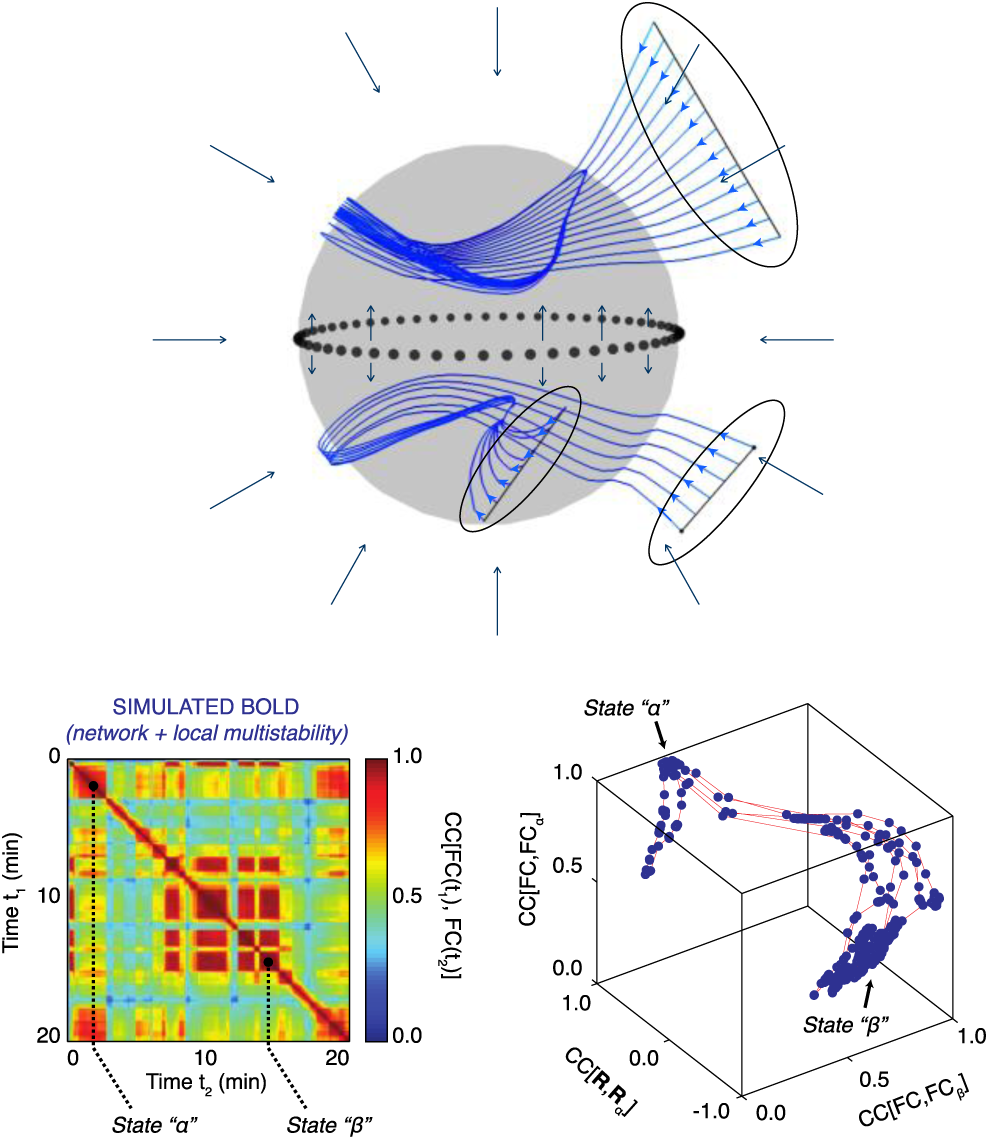
Structured Flows on Manifolds (SFMs). Upper figure shows a spherical attractive manifold, displaying various sets of initial conditions (black ovals) of trajectories (blue), evolving rapidly towards the manifold and then continue evolving on the manifold on a slower time scale. The time scale separation is evidenced by the large angle between trajectory and manifold (around 90 degrees). The flow on the manifold is split into two domains, one lower and one upper hemisphere, partitioned by a seperatrix (dotted line). The trajectories trace out lines on the manifold, following the flow (black arrows). The SFMs display a bi-stable organization with closed circular orbits on both hemispheres. The two lower figures show a similar organization, as captured by BOLD signals simulated with TheVirtualBrain (Hansen, Battaglia, Spiegler, Deco, & Jirsa, 2015; Sanz Leon et al., 2013). On the left, Functional Connectivity Dynamics (FCD) are shown over 20min, in which two large segments of invariant Functional Connectivity (FC) are identified as states alpha and beta. For both time windows, a principal component analysis was performed spanning state-characteristic subspaces by the leading principal components. When the BOLD signals were projected into the characteristic subspaces, the trajectory of the brain signal is unfolded, identifying the manifolds and trajectories of the corresponding states (figure on bottom right).

The link of SFM to flows and emergence can be conceptualized from considering a piece of music. The analogy of the “brain as a symphony” has been made by often, and is used to illustrate the fact that the emergence of function comes not from the action of a single brain area, but rather the coordination amongst all elements (unlike a symphony, however, in the brain there is no conductor). SFM theory provides a formal framework for these concepts of “brain as a symphony”. In a symphony, one can isolate the individual instruments to characterize their unique contribution, but it is difficult to appreciate its role in the symphony without considering the relation to other instruments. The statement: “The whole is greater than the sum of its parts.” is appropriate here for both the symphony and the brain.

We can further develop this analogy to build the intuition about SFM, particularly in the context of how different temporal flows (e.g., melody and piano lines in a simple song) interact in supporting the emergent behaviour (the whole song). The melody and harmony often move in *different time scales*. Each can be comprehended on their own, but in a well-composed song, the relation between the lines brings a richness that is not present in either alone (e.g., *aggregate property*). This fluctuation between the melody and harmony evolves throughout the song. It is common in classical pieces for the opening melody to be repeated as a motif, but over a slightly different piano line, which may completely change the mood of the piece.

In the brain, a parallel to the symphony analogy can be drawn. As the instruments in the orchestra and musical abilities of the artists define constraints upon the symphony to emerge, the anatomical connectivity and dynamic characteristic of the brain regions (network nodes) specify the rules for the evolution of dynamics. As we shall see below, this is far from a trivial constraint, as the anatomy helps define any spatial and temporal constraints for potential network configurations. For example, all things being equal, it is more likely that adjacent regions in occipital cortex will interact rather than occipital and frontal regions, simply because the occipital and frontal areas have few connections between them, and those that are connected indirectly at a long distance, imposing a longer time delay for transmission. Thus, anatomy establishes a deterministic architecture that prevents random manifolds and flows from occurring. This architecture, set atop the (nonlinear) dynamics of neurons and connected populations of neurons establishes the set of *motifs* that are available for the brain to combine in the coordination of behaviour (Sporns & Kotter, 2004). We can refer to these as *functional* modes to emphasize that they can be both actual and potential configurations.

The asymmetries in the brain’s space-time structure, set by the structural connectivity, establishes a potential for multi-scale actions (Deco et al., 2013). The multi-scale temporal character of these modes is founded on the fact that complex processes arise in an organism-environment context that inherently covers multiple scales. Armed with functional modes as essential building blocks, we propose additional dynamics (called operational signals) on time scales slower and faster than that of the modes. The slower process effectively binds functional modes together into sequences. More precisely, given functional mode emerges via a competition process to temporally dominate the functional dynamics, after which it destabilizes and gives way to another mode (Haken, 2006; Perdikis, Huys, & Jirsa, 2011). The transient dynamics between modes can be triggered either by ‘internal’ events (as in pre-constructed sequences) or by ‘external’ ones (such as perceptual events). Once engaged, the temporal attractivity of a modes guarantees functional robustness, whereas transitions between modes underlies flexibility for meaningful changes. Further variability in the function may arise via additional dynamics operating on times scales faster than (or similar to) that of the modes. Accordingly, brain function is organized in multilevel dynamical hierarchies.

The hierarchical architecture is central to effective information processing, where different temporal and spatial scales interact in moving the system through behavioural repertoires. Information provided to the brain system is meaningful if and only if it qualitatively changes the “state” that the brain occupies at that moment. If an incoming signal does not change the state, then the information was not meaningful, and the incoming signal could equally have not been present. Thus, while local dynamics may change dramatically, they may not have an appreciable effect on the trajectory of the network, and thus rather than change the flow to a new part of the manifold, may only result in a trivial variation in the current trajectory. If these local dynamics intersect with larger-scale dynamics at a critical point, this can establish a new trajectory for the system, either within an existing SFM or moving to a new SFM and hence a new emergent behaviour.

## 4. Mathematical Description of SFM

SFMs are the mathematical objects capturing the dynamic properties required from a system capable of the behavior we have discussed thus far. The system under consideration is high-dimensional with *N* degrees of freedom and highly nonlinear. In order to allow for this system to generate low-dimensional behavior, that is *M* dimensions with *M<<N*, there must be a mechanism in place, capable of directing trajectories in the high-dimensional space towards the *M*-dimensional sub-space. Mathematically this translates into two components, that are associated with different time scales: first, the low-dimensional attractor space contains a manifold *f(.*) and attracts all trajectories on a *fast* time scale; second, on the manifold a structured flow *g(.*) prescribes the dynamics on a *slow* time scale, where here slow is meant in comparison to the fast dynamics towards the attractor. For compactness and clarity, imagine the state of the system is described by the *N-*dimensional state vector *q(t*) at any given moment in time *t.* Then we split the full set of state variables into the components *u* and *s* where the variables in *u* define the *M* task-specific variables linked to emergent behavior in a low-dimensional subspace (the functional network) and the *N-M* variables in *s* define the remaining recruited degrees of freedom. Naturally, *N* is much greater than *M* and the manifold in the subspace of the variables *u* has to satisfy certain constraints to be locally stable, then all the dynamics is attracted thereto (Pillai & Jirsa, 2017).

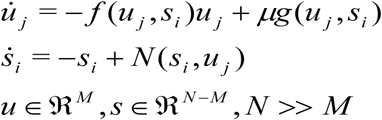

The flow of the nonlinear dynamic system is the right hand side of the differential equations. In state space, the flow is a form of force field that drives the state of the system along a trajectory. The tracing out of the trajectory is the evolution of the complex dynamic system, the flow is the rules that underlie the behavior. The above mathematical representation via a time-scale decomposition is not unique and there maybe other equivalent representations, capable of capturing the same flow in state space. However, the current representation is attractive for two reasons: 1) it provides a clear separation of the time scale via the smallness parameter *μ*, where the slow time scale is *μ*<<1 and the fast time scale is on the order of 1. 2) The current form has been successfully linked to networks composed of neural masses, coupled via multiplicative coupling functions, which are fundamental for the emergence of SFM (Pillai & Jirsa, 2017; Woodman & Jirsa, 2013). These multiplicative properties are at the heart of conductance-based modeling as embodied by the Hodgkin-Huxley equations, as well as essential in synaptic couplings. Mathematically the multiplicative coupling enables the manifold to be described globally, rather than only locally as it has been the case previously in formal theories of self-organization, such as Synergetics (Haken, 1996, 2006). The formulation of SFM is a general framework and the link to neuroscience is accomplished, for instance, when SFMs are derived from neural network equations. In these situations, the state vector *q(t*) is the vector of all activation variables across all brain regions and the SFM is the mathematical representation of the dynamics of the brain network. We will provide in the following examples of applications of SFM theory to neuroscience problems, which will in all cases refer to the state vector as neural activations. It is non-trivial and not lost on us, that the emergent SFM in brain activation space does not necessarily map isomorphically onto the low-dimensional dynamics (and thus SFM) in behavior. In other words, the lawfulness and rules underlying cognitive architectures may not be isomporphically related to the rules governing its directly associated brain dynamics. As attractive such isomorphism conceptually may be, it needs to be demonstrated empirically.

## 5. Modeling SFM in Epilepsy

A consideration about pathologies in the brain adds a critical element to our reflections on SFM and model emergence in the brain. Fundamental modeling of epilepsy has led to the postulate of the existence of a slow variable that dictates the expression of faster seizure activity (Jirsa, Stacey, Quilichini, Ivanov, & Bernard, 2014). During epileptic seizures, the firing activity of billions of neurons becomes organized so that oscillatory activity emerges that can be observed in electrographic recordings. This organization greatly reduces the degrees of freedom necessary to describe the observed activity, from single neurons firing to a few oscillatory collective variables. On the other hand, these oscillations trigger a series of processes at the microscopic level that slowly leads towards the end of the seizure. These slow processes can also be described by a collective variable, the permittivity variable that represents the balance (or imbalance) between the slowly varying pro- and anti-seizure mechanisms. *The fast variables span an SFM and the slow variable guides the brain system through the creation and annihilation of the SFM*. The composition of fast and slow variables in epilepsy is called the Epileptor.

Biophysical parameters that slowly change in the period preceding a seizure and during the ictal state are, for example, extracellular levels of ions (Heinemann, Konnerth, Pumain, & Wadman, 1986), oxygen (Suh, Ma, Zhao, Sharif, & Schwartz, 2006) and metabolism (Zhao et al., 2011). We can thus describe the evolution of a seizure with a few collective variables acting on different timescales: fast variables that, depending on the value of their parameter, can produce either resting or oscillatory activity with bifurcations separating the different regimes; slow variables describing the processes that brings the fast variables across the onset and offset bifurcations (Figure 2).

**Figure 2.**
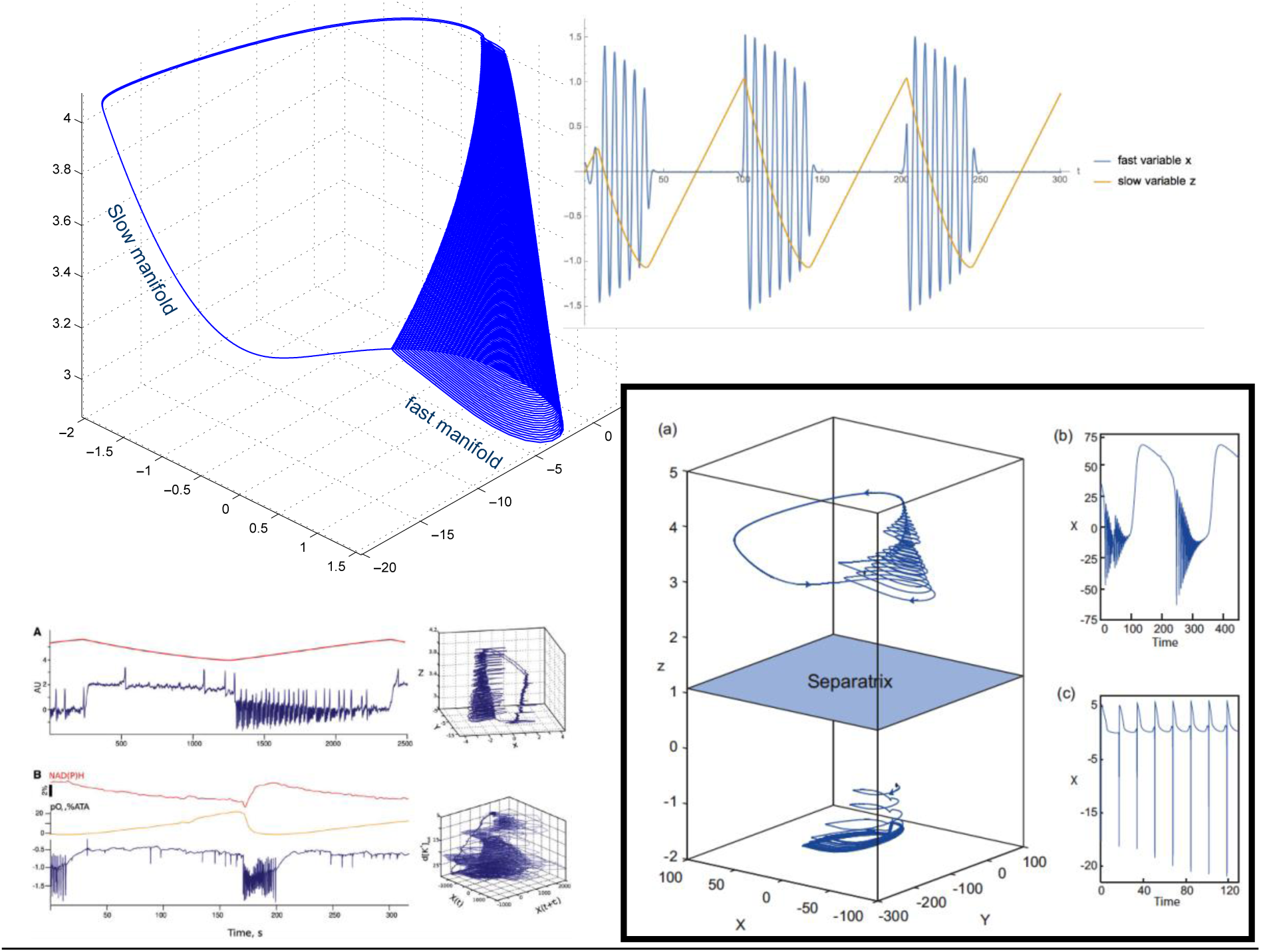
SFMs in Epilepsy. Ictal and non-ictal discharges have been characterized in nonlinear dynamics by two manifolds, a slow one-dimensional manifold illustrating the non-ictal resting state and a fast oscillation tracing out trajectories on a cone (see figure on the top). The corresponding time series are shown on the top-right, whereas the canonical SFMs are shown on the top-left. On the bottom left, two situations (time series, SFM) are shown for an empirical signal (B: rat, in-tuto hippocampus) and a detailed model signal (A: Epileptor), showing the identical topological features in state space. In the box on the bottom right, a state space is shown in (a), in which the upper region holds an Epileptor attractor and the lower region, separated by a separatrix (indicated in light blue), a so-far unknown attractor, which is hypothesized to be linked to refractory status epilepticus. The respective time series from the two attractor spaces are plotted in the two right panels (b) and (c)

Across multiple patients (Saggio, Spiegler, Bernard, & Jirsa, 2017), most had seizures characterized by different bifurcations in different moments, which implies that different classes of seizure types coexist and can be described with the same model, so that ultra-slow changes in the parameters of the fast variables can bring the patient closer to one or the other seizure type. From the perspective of dynamical system modeling, this states that there must exist some slow variable dynamics (under the assumption of autonomous systems). If the slow variable exists in pathological conditions, we make the assertion that slow variable dynamics plays an equally important role in healthy conditions evolving together with the fast variable dynamics as the actual emergent subsystem, or in Hermann Haken's words “order parameters”. The novelty here is that the emergent order parameters have an intrinsic time scale separation and comprise fast and slow variables, and not the typically single time scale of Synergetics. Fast variables act on slow variables and vice versa. The mutual presence of multiple time scales in the emergent system, the SFM, is reflective of the adaptive nature of the brain.

## 6. Hidden capacity of networks revealed in the SFM framework

We noted earlier that a distinct advantage of creating models of rules governing coordination of behavior is the possibility of identifying novel configurations that had not yet been expressed or observed. This advantage can be illustrated from further elaboration of the Epileptor model. A parameter sweep of the model provides confirmation of the interplay of fast and slow variables in moving the system from a quiescent phase, into seizure, and then back out. A broader parameter search identified another SFM, in which the system engaged in broad slow oscillations (Figure 2, bottom right) (El Houssaini, Ivanov, Bernard, & Jirsa, 2015). Phenomenologically, these trajectories resembled what is seen in refractory status epilepticus (RSE). The critical aspect of this observation was that this repertoire was not obvious in the initial creation of the model, but this “new behavior” was in fact part of the lawful behavior of the system.

The second important aspect of this was the observed dependencies of the seizure and RSE behaviors, wherein modification of slow variables allowed a transition between behavior, which was confirmed in animal models (El Houssaini et al., 2015). This is also a vital observation clinically as it suggests a different treatment path to alleviating RSE is to re-establish seizure rather than eliminate the dynamics all together.

By modeling the system, rather than a given realization, we were able to identify this hidden state that would be invisible to other approaches that attempt only to characterize the timeseries/realizations. As we noted earlier, even if one captures a large number of realizations, the quantification of these only is relevant to the particular behavior and not to the function of the system. Modeling the system, similar to what we propose in with SFM, captures both what the system does when you are watching and what is could do you when you are not. The Epileptor perfectly embodies this where the model captured the presence of the RSE state, even though the system did not need to generate a realization to know that the state existed.

The Epileptor model gives a very salient demonstration of the use of SFM framework to under “disease potential”. This yields from two postulates. The first stems from the physiological fact that anyone’s brain has the potential to show seizures given the right conditions. From the SFM perspective, what this suggests is that “seizure” is an existing repertoire in anyone’s brain that can be expressed when the parameters are right (Jirsa et al., 2014). The phenomenological model provides a useful characterization of the state changes that need to occur in order to shift the flows on the manifold to the seizure attractor. A further exploration of the Epileptor model indicated that another behaviour can be expressed, namely that of RSE, again once the control parameter changes are sufficient to move from the seizure attractor to the RSE attractor.

The second postulate stems from the first. If epilepsy is a part of the natural repertoire of the brain, can other clinical conditions be similarly regarded? At face value, the suggestion would be ‘no’ because epilepsy may be in inherent biophysically property of oscillatory networks, while other scenarios arising from acquired brain injury or neurodegenerative disorders may not be equally represented across the population. But perhaps we can recast the perspective somewhat. As an adaptive system, the brain is in a constant state of testing new configurations to enhance capacity (Minerbi et al., 2009; Ziv & Brenner, 2018). While this is generically considered plasticity, these reconfigurations seem to be spontaneous and persist when the outcome is adaptive, which is property of complex adaptive systems. These changes are considered atop a more stable repertoire, which, usually, prevents a catastrophic situation where a maladaptive configuration is reinforced. This gives us a segue to consider maladaptive responses in terms of clinical outcome. It may be the case that brain disease expression reflects maladaptation. This would explain the observations where two persons with ostensibly the same damage can show markedly different clinical expressions, one showing severe impairment and the other showing much less, if any. In the first case, there is an attempt to adapt the damage but the new manifold or attractor that emerged was maladaptive, resulting a dysfunctional realization. In the second case, the adaptation was more robust allowing the person’s to maintain more stable manifold, reducing the clinical severity. Thus, unlike the epilepsy case where seizure is a naturally part of the brain’s repertoire, in other cases the clinical expression is reflected a given brain’s capacity to adapt to a pathological process.

These two postulates can be unified under the idea that the facility with which one moves from one manifold to another will dictate clinical outcome. For epilepsy, many persons will never have a seizure, suggesting the despite the existence of the seizure manifold, the system configuration is such that moving to this manifold never happens. In the case of perturbation from acquired brain injury or degenerative disorders, the maladaptive response comes because the existing system repertoire was not able to accommodate the perturbation. Where the clinical outcome is less severe, the perturbation still has a negative effect, but the existing repertoire is able to adapt sufficiently so as to limit disability.

The perspective changes the way we consider clinical progression from one where the brain is static and the clinical expression is simple the loss of function to one where the clinical progression is an expression of the continual adaptation of the brain. The adaptation itself may indeed be as debilitating as the triggering event.

If this is true, then it should be possible to characterize the capacity of a given brain to adapt to negative perturbation by construction and exploration of a person’s SFM. An even more intriguing potential is that such a characterization may suggest a course of intervention that makes use of the capacity of a given person to traverse their SFM and adapt.

## 7. Future directions and final thoughts

There already exists a growing body of work that characterizes neurophysiological data using dimensionality reduction techniques that is a step towards defining low-dimensional manifolds that constrain network flows (Gallego et al., 2018). Indeed, recent work in functional neuroimaging is focusing on the configurations of functional networks and the changes in their configurations in relation to behavior (Khambhati, Sizemore, Betzel, & Bassett, 2018; Shine et al., 2019). Analysis of the changes in functional networks per se, also referring to a functional connectivity dynamics (Figure 1) (Hansen et al., 2015; Hutchison et al., 2013), provides a relatively straight path to manifold estimation.

There are established methods for manifold estimation that extend beyond functional connectivity and instead define the space (i.e., manifold) that constrains the variance of specific neurophysiological signals. Here, trial-by-trial signals are considered together to define the dimensionality of the system and then characterize the manifold features (Gallego, Perich, Miller, & Solla, 2017). Methods such as principal components analysis can give access to the manifold space (Banerjee, Tognoli, Assisi, Kelso, & Jirsa, 2008), while others explicitly characterize the manifold such as Stochastic Neighborhood Embedding and Uniform Manifold Approximation and Projection (Ma & Fu, 2011). Algebraic Topology methods are also proving to be powerful complementary techniques by giving access to geometrical characterizations of manifolds that can then be related to cognition and behavior. For example, Saggar and colleagues looked at topological structures in relation to cognitive performance fMRI data (2018), finding that those with a more distributed topology showed better cognition (Figure 3). Additional features of estimated manifolds, such as switching, dwell time, transitional probabilities, are important aspects that emphasize the temporal flows on the manifold. Along these lines, an emphasis on trial-by-trial time series, rather than simple averages of data, are preferable. Differences in average features may have some utility in selection of key nodes for network identification, but obliterate the higher order statistical moments of the data, which are central to SFM expression.

**Figure 3.**
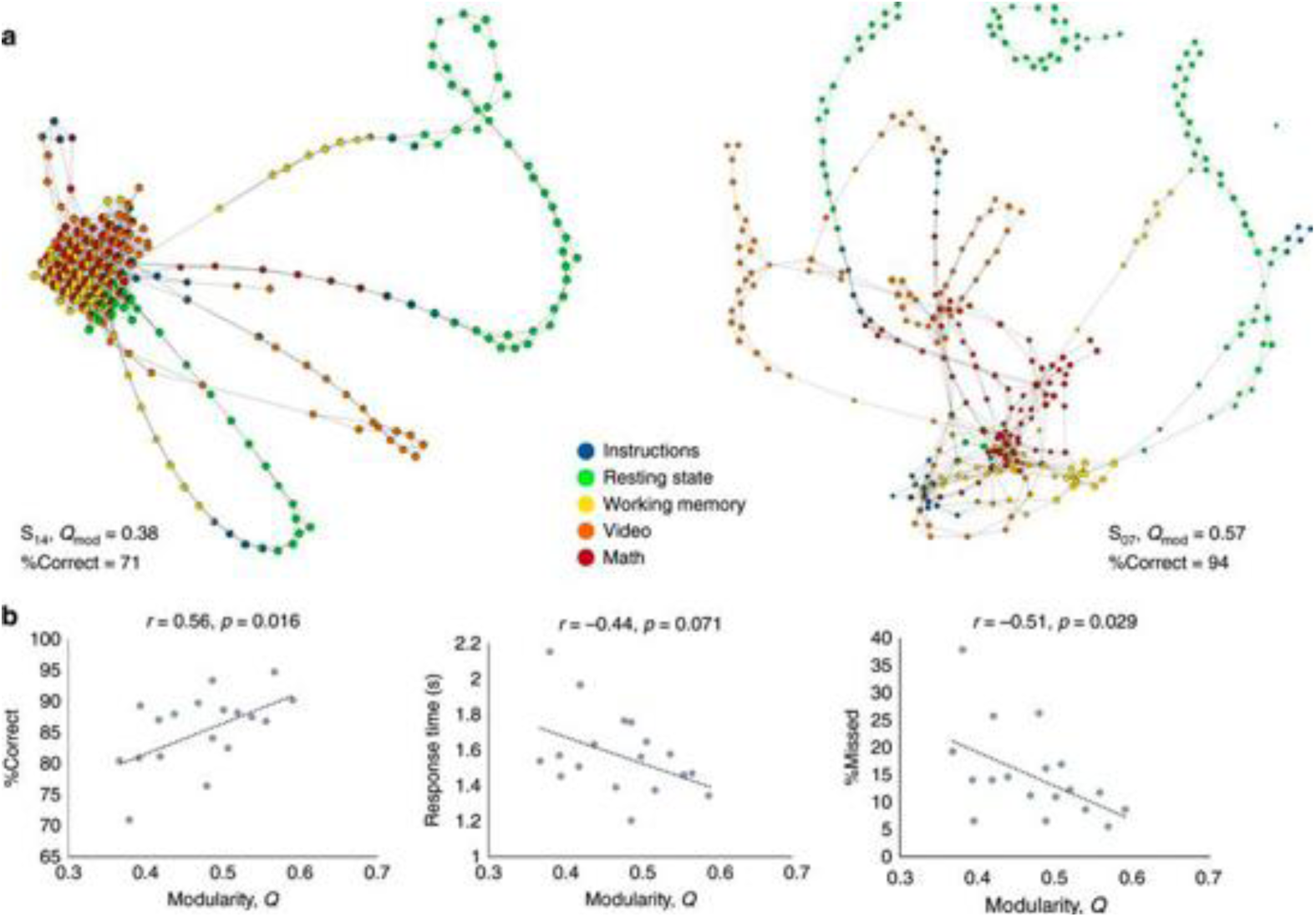
Excerpt from Saggar et al (2018) showing the comparison of shape graphs constructed using Topological Data Analysis of the dynamics of network evolution measured with fMRI. Graphs for two subjects (top panel a) are shown and were quantified based on modularity, with S14 (left) showing low modularity and S07 (right) showing higher modularity. Panel b shows the correlation between modularity indices across all subjects and different aspect of behavior. The pattern suggests subject with higher modularity, which may suggest of a more complex manifold architecture, have better behavior.

There is an additional aspect that highlights the unique aspect of the SFM framework, which is that the behavior that emerges from the brain must also be characterized as flows on manifolds. This enables a new level of analysis to better characterize brain-behavior relationship in terms how the specific evolution of flows on manifolds in brain constrain and are constrained by the flows on manifolds in behavior. Here there are fewer methods that map such interdependency between flows, though some candidates do exist (Breakspear & Terry, 2002; Flack, 2017; Terry & Breakspear, 2003).

This begs the question as to whether cognitive processes, such as memory and emotion can be characterized under the SFM framework. Although most behavioral measures of cognition are often single points, such as reaction time or accuracy of responses, the notion of mental flows is pervasive in theory (Spivey, 2007). A recent expression emphasizes a seamless flow capturing the process of moving between sensation and action and back where the lines between traditional states (e.g., sensation, perception, memory) is blurred if not absent. For cognitive processes this is a challenge as they are not easily measured. However, their impact on ongoing behavior, such as eye movements or reaching (Song & Nakayama, 2009), has been used successfully to characterize the dynamics of processes and does give a potential access point for the creation of behavioral SFMs that can be linked to brain SFMs. The trajectories create a personal space that can be translated to a manifold. Across realizations, the flows along the manifold can then be related with the corresponding flow elicited in the brain – essentially mapping SFMs in behavior to those of the brain. This will yield new understanding of how the richness of behavior that we observe is enabled by the richness brain dynamics that we measure.

## Acknowledgements

This work was supported by a Canadian NSERC grant RGPIN-2018-04457 (A.R.M.) and European Union’s Horizon 2020 research and innovation program grant 720270. (V.K.J.)

## Author Contributions

A.R.M. and V.K.J. contributed equally to this paper.

